# Microtiter plate-based antibody-competition assay to determine binding affinities and plasma/blood stability of CXCR4 ligands

**DOI:** 10.1101/2020.07.25.221085

**Authors:** Mirja Harms, Andrea Gilg, Ludger Ständker, Ambros J. Beer, Benjamin Mayer, Volker Rasche, Christian W. Gruber, Jan Münch

## Abstract

C-X-C chemokine receptor type 4 (CXCR4) is involved in several intractable disease processes, including HIV infection, cancer cell metastasis, leukemia cell progression, rheumatoid arthritis, asthma and pulmonary fibrosis. Thus, CXCR4 represents a promising drug target and several CXCR4 antagonizing agents are in preclinical or clinical development. Important parameters in drug lead evaluation are determination of binding affinities to the receptor and assessment of their stability and activity in plasma or blood of animals and humans. Here, we designed a microtiter plate-based CXCR4 antibody competition assay that enables to measure inhibitory concentrations (IC_50_ values) and affinity constants (K_i_ values) of CXCR4 targeting drugs. The assay is based on the observation that most if not all CXCR4 antagonists compete with binding of the fluorescence-tagged CXCR4 antibody 12G5 to the receptor. We demonstrate that this antibody-competition assay allows a convenient and cheap determination of binding affinities of various CXCR4 antagonists in living cells within just 3 hours. Moreover, the assay can be performed in the presence of high concentrations of physiologically relevant body fluids, and thus is a useful readout to evaluate stability (i.e. half-life) of CXCR4 ligands in serum/plasma, and even whole human and mouse blood *ex vivo*. Thus, this optimized 12G5 antibody competition assay allows a robust and convenient determination and calculation of various important pharmacological parameters of CXCR4 receptor-drug interaction and may not only foster future drug development but also animal welfare by reducing the number of experimental animals.

## Introduction

The G protein-coupled receptor CXCR4 (CXC chemokine receptor 4) is a 352 amino acid cell surface protein with CXCL12 as its sole endogenous chemokine ligand^1,2,3^. The CXCR4/CXCL12 pair plays an important and unique role in cellular trafficking processes involved in organ development, hematopoiesis, vascularization, in cell and tissue renewal and regeneration, in inflammation and immune control, stem cell homing and mobilization^4,5,6,7,8^. Aberrant CXCR4/CXCL12 signaling is associated with a variety of pathophysiological conditions including cancer metastasis, enhanced tumor growth, chronic inflammation, leukemia and altered immune responses^9^. Furthermore, CXCR4 is a major coreceptor of HIV-1 during the late stages of infection and associated with rapid disease progression^10,11^. Thus, CXCR4 represents an important drug target. Several small molecules (e.g. AMD3100^12^, AMD070^13^), peptides (e.g. Polyphemusin 2^14^ or Ly2510924^15^), or nano- and antibodies (e.g. Ulocuplumab^16^ or ALX-0651^17^) that target and antagonize CXCR4 have been identified^9^ and represent promising leads for drug development. Diverse animal studies provided evidence that CXCR4 inhibitors allow mobilization of hematopoietic stem cells, suppression of CXCR4-tropic HIV-1 replication and reduction in tumor growth and/or metastasis. Several clinical studies that evaluate CXCR4 antagonists as therapeutic agents in e.g. pancreatic cancer, adenocarcinomas, multiple myeloma or myelokathexis are currently running. However, none of these compounds has been approved for the treatment of chronic diseases like cancer or HIV/AIDS. The main reason is likely because long-term inactivation of CXCR4 also inhibits the physiological function of the receptor and causes side effects^18^. The only licensed CXCR4 antagonist to date is AMD3100 (Plerixafor), which is used as single injection to mobilize hematopoietic stem cells in cancer patients^9^. AMD3100 is a bicyclam small molecule drug that was initially developed as a treatment against CXCR4-tropic HIV-1 infection but failed during long term application studies^18,19^. In the last years, several analogues of AMD3100 were developed covering a range of sub-structural types. One of them is the non-cyclam small molecule AMD070 that is orally bioavailable and was revived in several clinical trials including a phase III clinical trial for WHIM patients^22,21^. Another orally available small molecule is MSX-122 that was described as a partial antagonist of CXCR4 and is currently investigated in a phase II clinical trial as an oral drug for hot flashes in breast cancer-positive post-menopausal women^20,21^. Two other orally available small drug candidates are Burixafor (TG-0054) that is a monocyclic CXCR4 antagonist und currently tested in a phase II study for stem cell mobilization^23,21^, and the isothiourea compound IT1t that was used for crystallization of the CXCR4 receptor thereby revealing a distinct binding mode from AMD3100 within the receptor binding pocket^24,25,26^.

Besides small molecules, several peptide-based CXCR4 antagonist are also being tested for multiple applications. Those peptides are often based on naturally occurring ligands, e.g. the 17 amino acid CXCL12 analogue CTCE-9908^27^, and the two cyclic peptides LY2510924^15^ and BKT-140^28^ that are currently tested in phase II clinical trials determining their effect on different kinds of cancer and stem cell mobilization^21,29 (for a review see 30)^. Another linear peptide drug candidate is EPI-X4 (Endogenous Peptide Inhibitor of CXCR4), a recently identified body own antagonist of CXCR4^31,32,33^. This 16-mer peptide encompasses residues 408-423 of human serum albumin and is proteolytically released from its precursor by acidic aspartic proteases^34^. EPI-X4 blocks CXCL12-mediated signaling, thereby preventing the migration of cancer cells, mobilizing hematopoietic cells and inhibiting inflammatory responses *in vitro* and in mouse models. In addition, EPI-X4 is also an effective inhibitor of CXCR4-tropic HIV-1. This endogenous peptide represents an interesting lead for further clinical development because it is not immunogenic and, in contrast to AMD3100, does not bind CXCR7^31,35^. In addition, it also acts as inverse agonist and reduces the basal signaling activity of CXCR4, and does not exert mitochondrial cytotoxicity. However, the short half-life of the peptide in plasma (t_1/2_ ~ 17 min)^31^ and its only median potency in antagonizing CXCR4 (IC_50_ to suppress CXCL12 migration in median micromolar range) prevent its further clinical development as CXCR4 targeting drug, at least for systematic therapy. A structure–activity relationship study allowed to reduce the size and to improve the CXCR4-antagonizing potency of EPI-X4^31,36^. One example of an optimized EPI-X4 derivative is WSC02 that encompasses only 12 residues and inhibits CXCR4-tropic HIV-1 infection and suppresses leukemia cell migration towards CXCL12 with IC_50_ values in the nanomolar range. However, despite the improved CXCR4 antagonizing potency, EPI-X4 and its first-generation derivatives suffer from a low stability in plasma due to proteolytic inactivation of the peptide by leucyl aminopeptidases^31,34^.

Proteolytic degradation often limits systemic therapeutic applications of peptide-based drugs. Increasing peptide stability relies often on evaluating large numbers of lead compounds, which requires suitable *in vitro* techniques providing reliable predictions of *in vivo* performance and reducing the number of animal experiments^37^. Peptide stability and proteolysis is usually determined *ex vivo* by spiking the peptide into plasma, serum or whole blood (derived from humans or mice), incubation at 37°C and subsequent analysis of the specimen by means of chromatography and/or mass spectrometry^37,38,39^. However, these techniques are time consuming, do not permit high throughput testing and are not predictable of *in vivo* stability, which is of particular interest for peptide-based drugs because of the ease in synthesizing modified variants, as compared to small molecules or antibodies. We here set out to design a microtiter plate-based screening assay that aims to determine the stability of CXCR4 ligands in plasma or serum as well as whole blood. We took advantage of the fact that most if not all CXCR4 antagonists block CXCL12 binding by occupying a region formed by the second extracellular loop (ECL2) of CXCR4. ECL2 is also the binding site of the anti-CXCR4 antibody 12G5^40,41^. Thus, most CXCR4 antagonists compete with binding of 12G5 to CXCR4-expressing cells which can be measured and quantified by flow cytometry. We demonstrate that this assay allows a robust and convenient determination of ligand-receptor pharmacodynamics and to measure the stability of CXCR4 ligands in serum/plasma and even whole human and mouse blood.

## Results

### CXCR4 antagonists compete with 12G5 for binding to CXCR4

It has previously been demonstrated that several CXCR4 antagonists, including AMD3100^42,43^, EPI-X4^31^, T140^44^, T134^45^, ALX40-4C^46^, and POL3026^47^ compete with the monoclonal anti-CXCR4 12G5 antibody for binding to CXCR4. To corroborate and expand these findings, several small molecules and peptide-based CXCR4 antagonists (Table 1) were analyzed for competition with 12G5. As shown in Fig. 1a, all small molecules (with the exception of MSX-122) resulted in a concentration-dependent reduction of 12G5 binding. IT1t and Burixafor competed most efficiently with 12G5, with IC_50_ values of 1.7 and 0.3 nM, respectively, whereas AMD070 and AMD3100 were slightly less active (IC_50_ values of 37 and 578 nM, respectively) (Table 2). MSX-122, a partial CXCR4 antagonist did not interfere with 12G5 binding, which is likely due to the fact that it does not bind to ECL2 but rather penetrates into the deep binding pocket in CXCR4^20^. In addition, all peptidic antagonists analyzed reduced 12G5 antibody binding in a concentration-dependent manner. Here, the optimized EPI-X4 derivate WSC02, the respective dimeric analog WSC02×2, and LY2510924 were most active (IC_50_ values of 290, 182 and 151 nM, respectively), followed by BTK-140, endogenous EPI-X4 and CTCE9908 (Fig. 1b, Table 2). None of the compounds interfered with binding of the 1D9 antibody, which recognizes the N-terminal flexible domain of CXCR4 (Fig. S1a and b). BTK140 seemed to reduce 1D9 binding, but this could be attributed to cytotoxic effects caused by this compound at concentrations above 10 μM (Fig. S1c). Thus, all assayed CXCR4 ligands (except the partial antagonist MSX-122) prevent binding of the 12G5 antibody.

**Table 1.**
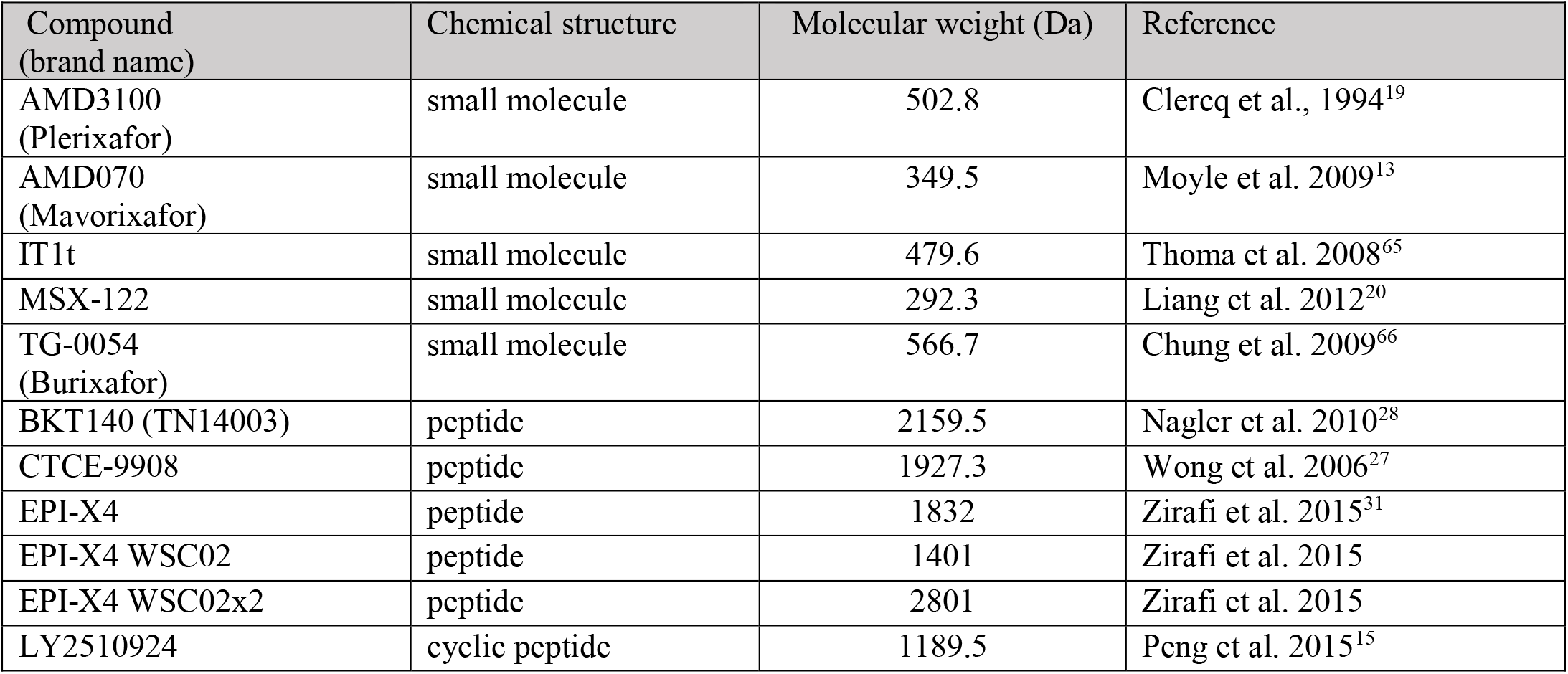
List of the CXCR4 antagonists used in this study.

| Compound (brand name) | Chemical structure | Molecular weight (Da) | Reference |
| --- | --- | --- | --- |
| AMD3100 (Plerixafor) | small molecule | 502.8 | Clercq et al., 1994<sup>19</sup> |
| AMD070 (Mavorixafor) | small molecule | 349.5 | Moyle et al. 2009<sup>13</sup> |
| IT1t | small molecule | 479.6 | Thoma et al. 2008<sup>65</sup> |
| MSX-122 | small molecule | 292.3 | Liang et al. 2012<sup>20</sup> |
| TG-0054 (Burixafor) | small molecule | 566.7 | Chung et al. 2009<sup>66</sup> |
| BKT140 (TN14003) | peptide | 2159.5 | Nagler et al. 2010<sup>28</sup> |
| CTCE-9908 | peptide | 1927.3 | Wong et al. 2006<sup>27</sup> |
| EPI-X4 | peptide | 1832 | Zirafi et al. 2015<sup>31</sup> |
| EPI-X4 WSC02 | peptide | 1401 | Zirafi et al. 2015 |
| EPI-X4 WSC02x2 | peptide | 2801 | Zirafi et al. 2015 |
| LY2510924 | cyclic peptide | 1189.5 | Peng et al. 2015<sup>15</sup> |

**Table 2.** Anti-CXCR4 activity and K_i_ values of the CXCR4 ligands.

| Compound | IC<sub>50</sub> 12G5 (nM)<sup>1</sup> | IC<sub>50</sub> HIV-1 (nM)<sup>2</sup> | IC<sub>50</sub> Migration (nM)<sup>3</sup> | K<sub>i</sub> (nM)<sup>4</sup> |
| --- | --- | --- | --- | --- |
| AMD3100 | 578 ± 105 | 36 ± 6 | 60 ± 13 | 221 ± 40 |
| AMD070 | 37 ± 15 | 23 ± 4 | 152 ± 3 | 14 ± 6 |
| IT1t | 1.5 ± 1 | 5 ± 0.9 | 230 ± 20 | 0.6 ± 0.4 |
| MSX-122 | >100,000 | >10,000 | Nd | - |
| TG-0054 | 0.3 ± 0.04 | 78 ± 6 | 1401 ± 259 | 0.1 ± 0.02 |
| BKT-140 | 508 ± 308 | 435 ± 153 | 932 ± 131 | 194 ± 117 |
| CTCE-9908 | 16,205 ± 3626 | - | 6912 ± 2606 | 6,185 ± 1384 |
| EPI-X4 | 1,775 ± 162 | 31,390 ± 3288 | Nd | 677 ± 62 |
| EPI-X4 WSC02 | 290 ± 75 | 257 ± 109 | 15,824 ± 3502 | 111 ± 29 |
| EPI-X4 WSC02x2 | 182 ± 31 | 254 ± 103 | 2,672 ± 233 | 69 ± 12 |
| LY2510924 | 151 ± 78 | 8 ± 0.8 | 66 ± 8 | 58 ± 30 |

<sup>1</sup> IC<sub>50</sub> shown are means +/- SEM derived from 3 independent 12G5 antibody competition assays. <sup>2</sup> IC<sub>50</sub> shown are mean +/- SEM values obtained from three independent HIV-1 infection assay each performed in triplicates. <sup>3</sup> IC<sub>50</sub> shown are mean +/- SEM values obtained from three independent migration assays each performed in triplicates. <sup>4</sup> K<sub>i</sub> values were calculated as described in the text. Nd = not determined

**Figure 1.**
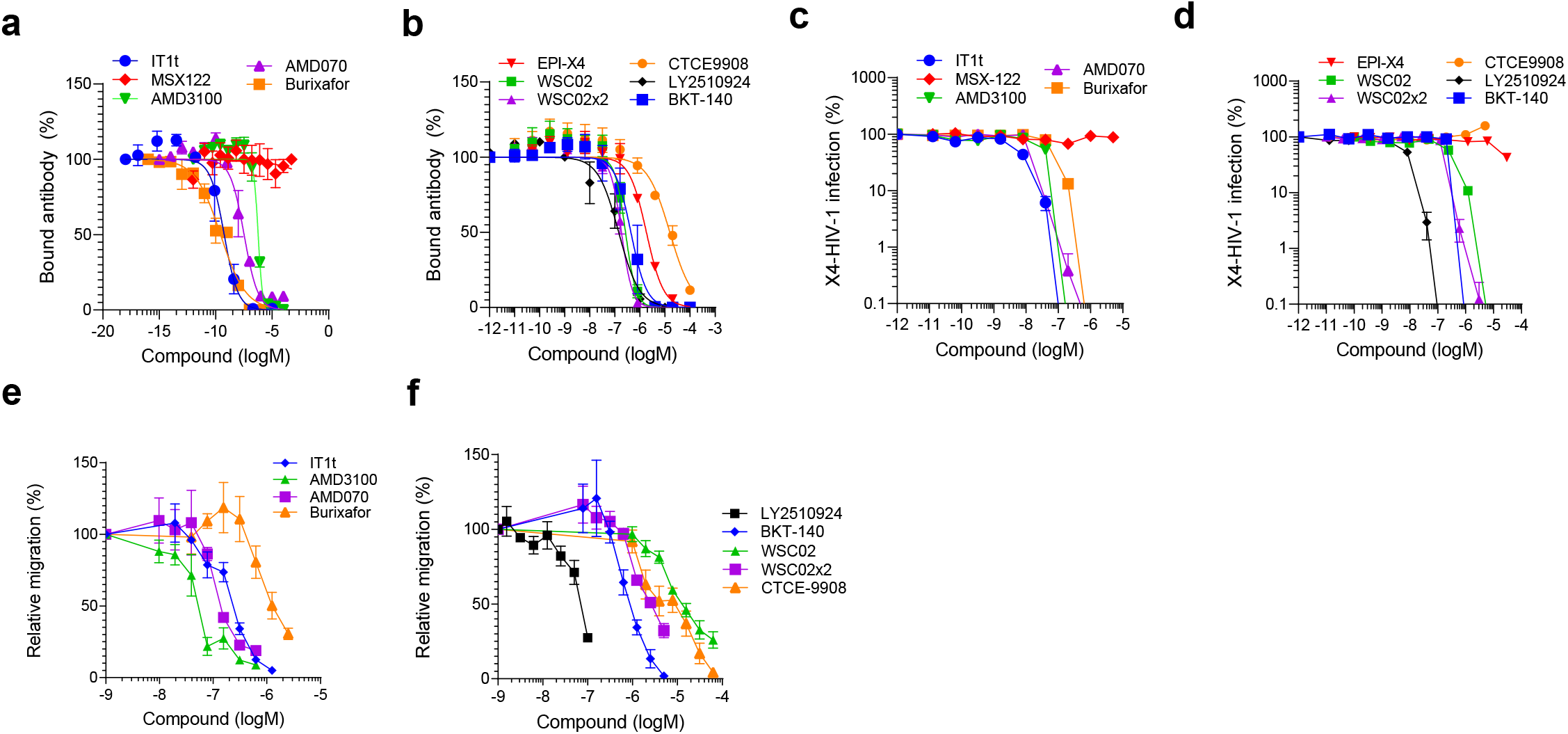
CXCR4 antibody competition, inhibition of HIV-1 infection, and suppression of T cell migration by CXCR4 ligands. (a, b) Competition of small molecule (a) and peptide (b) ligands with CXCR4 ECL-2 specific 12G5 antibody. Small molecules and peptides were diluted in PBS and added to SupT1 cells. A constant concentration of APC-labelled 12G5-antibody was added immediately afterwards. After 2 hours incubation in the dark, cells were washed and analyzed by flow cytometry. Data shown are average values derives from 3 individual experiments ± SEM. (c, d) Inhibition of HIV-1 infection by CXCR4 ligands. TZM-bl cells were incubated with serial dilutions of compounds for 15 min before they were inoculated with CXCR4-tropic HIV-1. Infection rates were determined after 3 days by β-galactosidase assay. Data shown were derived from 3 individual experiments performed in triplicates ± SEM. (e, f) Migration of SupT1 cells in the presence of CXCR4 ligands towards a CXCL12 gradient. The assay was performed in a transwell plate with 5 μm pore size with 100 ng/mL CXCL12 in the lower chamber. The number of migrated cells was determined by CellTiterGlo® assay. Shown are average values derived from 3 individual experiments performed in triplicates ± SEM.

### CXCR4 antagonists inhibit CXCR4-tropic HIV-1 infection and block CXCL12-dependent cell migration

To analyze the CXCR4 ligands in functional assays, we determined their activity against CXCR4 tropic HIV-1 in TZM-bl reporter cells. All small molecule CXCR4 ligands (except MSX-122) suppressed HIV-1 infection with IC_50_ values in the range of 6 nM for IT1t and 77 nM for Burixafor (Fig. 1c, Table 2). Similarly, all peptide antagonists (except for CTCE-9908) blocked viral infection (Fig. 1d, Table 2), with LY25110924 being the most potent inhibitor (IC_50_ of 8 nM), followed by WSC02×2 (204 nM) and BTK-140 (293 nM), WSC02 (279 nM) and endogenous EPI-X4 (31 μM). Finally, we determined the IC_50_ of the CXCR4 ligands in antagonizing SupT1 T cell migration towards CXCL12. All tested CXCR4 ligands suppressed migration (except MSX-122). For small molecules we obtained IC_50_ values between 60 nM (AMD3100) and 1400 nM (Burixafor) (Fig. 1e, Table 2), and for peptides 66 nM for LY2510924, and 15 μM for WSC02 (Fig. 1f, Table 2). Thus, with the exception of MSX122 and CTCE-9908, all analyzed CXCR4 ligands bind CXCR4, prevent 12G5-antibody binding, CXCL12-induced cellular migration and X4-tropic HIV-1 infection.

### Determination of K_i_ values for CXCR4 ligands

The Cheng-Prusoff transformation (see formula 3 in material/methods section)^48^ was used for calculation of equilibrium inhibition constants (K_i_) from IC_50_ values. To ensure that an equilibrium between ligand and antibody is established under the experimental conditions of the antibody competition assays, we first determined IC_50_, IC_90_ and areas under the curves (AUC) values for EPI-X4 WSC02 after different incubation times. No significant differences were obtained for these values after 1, 2 and 4 hours, showing that an equilibrium was established in the competition assay (Fig. S2). Consequently, all subsequent experiments were performed with a 2-hour incubation period. Since the Cheng-Prusoff transformation requires knowledge of the dissociation constant (K_d_) of the competing ligand, we experimentally determined the K_d_ of the antibody by establishing a saturation binding curve under equilibrium conditions (for reference see ^49, 25^). For this, a defined number of 5,000 SupT1 cells per well was incubated with increasing concentrations of the labelled 12G5 antibody, until saturation was reached (B_max_) (Fig. 2a). To exclude unspecific binding, the same experiment was performed in the presence of a high concentration of an optimized EPI-X4 variant (2000 μM, >10,000-fold its IC_50_) (Fig. 2a). The unspecific binding signal was then subtracted to obtain values for the specific binding curve. Fig. 2b shows the specific binding curve summarized for 4 individual titrations. From this curve, the half maximal binding concentration [Ab]_0.5_ of the antibody was determined as 155.5 pM (Fig. 2b, Table S1). Since the K_d_ is dependent on the total receptor concentration (R_t_) used for the reaction, we next quantified surface CXCR4 per cell using quantitative flow cytometry. SupT1 cells were incubated with the 12G5 mAb in saturation. Afterwards cells were stained with labelled secondary antibodies and simultaneously with FACS beads harboring known mAb concentrations. Mean fluorescence (MFI) signals of the beads (Fig. 2c) were used to define a standard curve (Fig. 2d) with that the MFI signal of the cells was interpolated. This approach allowed to calculate in average ~ 29,000 CXCR4 molecules per SupT1 cell, that fit with published data^50^. Based on the total receptor concentration in the assay we then applied formulas 1 and 2 (see methods) to calculate the K_d_ of the 12G5 mAb with 151.5 pM (Table S1). Notably, IC_50_ values obtained for CXCR4 antagonists in competition assay performed with a 10-fold increased number of cells (5 ×10^4^) did not change significantly, suggesting that the assay yields robust data even over a broad range of cell numbers (Fig. S3). Next, K_i_ values were calculated for all CXCR4 ligands according to formula 3 (Table 2), based on the IC_50_ values obtained in the competition assays, revealing e.g. sub-nanomolar K_i_ values for the small molecules TG-0054 (0.1 nM) and IT1t (0.6 nM) and mostly nanomolar K_i_ values for the remaining CXCR4 ligands (Table 2). Thus, the CXCR4 competition assay can be conveniently performed in a microtiter format and allows rapid and quantitative determination of ligand receptor pharmacodynamics.

### Adaption of the competition assay to determine plasma stability of CXCR4 ligands

We were wondering whether this assay may also allow determining the stability of CXCR4 ligands, in particular peptides, in human blood plasma. First, we analyzed whether plasma alone (pooled from 6 donors) interferes with 12G5 binding to CXCR4. As shown in Fig. 3a, a cell culture concentration of 50 % plasma did not affect antibody binding. Next, we studied proteolytic inactivation of the EPI-X4 WSC02 derivative in plasma. For this, the peptide was diluted in 99.3 % plasma to 20 μM and mixtures were agitated at 37°C. At different time points (minutes to hours), aliquots were taken, serially diluted and then analyzed in the competition assay (Fig. 3b). We observed a concentration-dependent and complete suppression of 12G5 binding by WSC02 samples that were taken and analyzed at t = 0 min. Samples obtained at later time points gradually lost their “function” of blocking antibody binding, which is likely due to proteolytic inactivation of the peptide in plasma (Fig. 3b). In fact, addition of L-leucinethiol, an inhibitor of leucyl aminopeptidases that inactivate EPI-X4^31,34^, largely prevented WSC02 inactivation in plasma (Fig. S4). Of note, even after 24 hours of incubation, WSC02 remained partially active and competed with 12G5 binding.

**Figure 2.**
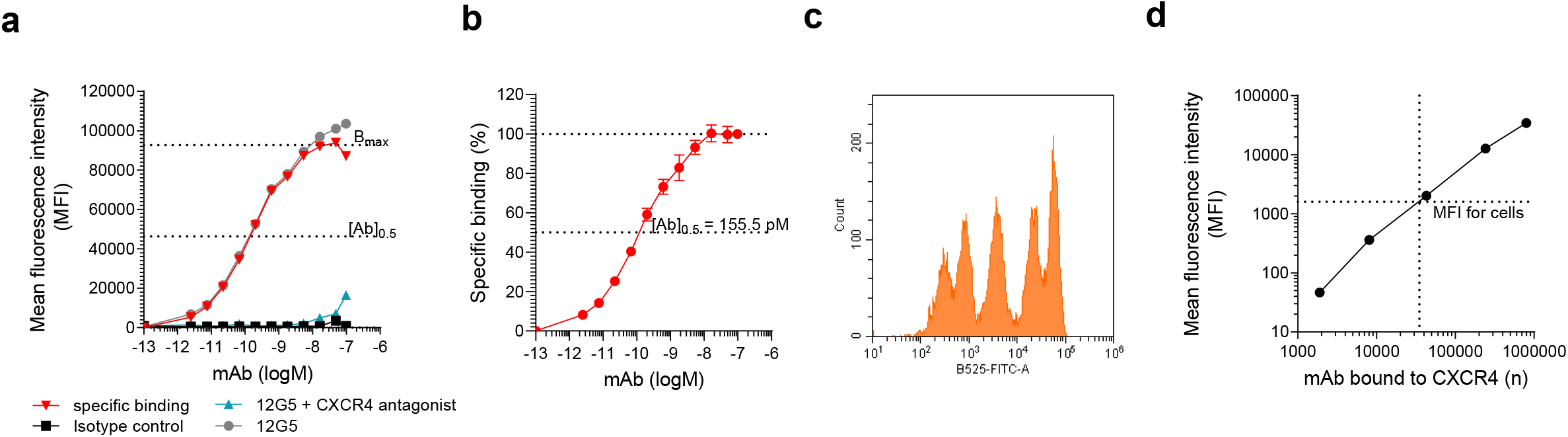
Determination of the dissociation constant (Kd) of the 12G5-antibody using flow cytometry. (a) The 12G5-APC antibody was titrated on SupT1 cells. For determination of unspecific binding a CXCR4 antagonist (derivative of WSC02) was applied at its 10,000 fold K_d_ during antibody binding and subtracted from the 12G5-binding curve to determine specific antibody binding. Data shown are from one representative experiment. (b) Averaged specific binding curve derived from 4 individual experiments ± SEM. (c, d) CXCR4 concentration on SupT1 cells was determined using quantitative flow cytometry. The MFI of SupT1 cells labelled with a primary anti-CXCR4 mouse monoclonal antibody (mAb) and a secondary anti-mouse antibody (c) was compared to a calibration curve derived from bead populations harboring a specific amount of mAb at their surface (d). Shown is one representative experiment.

**Figure 3.**
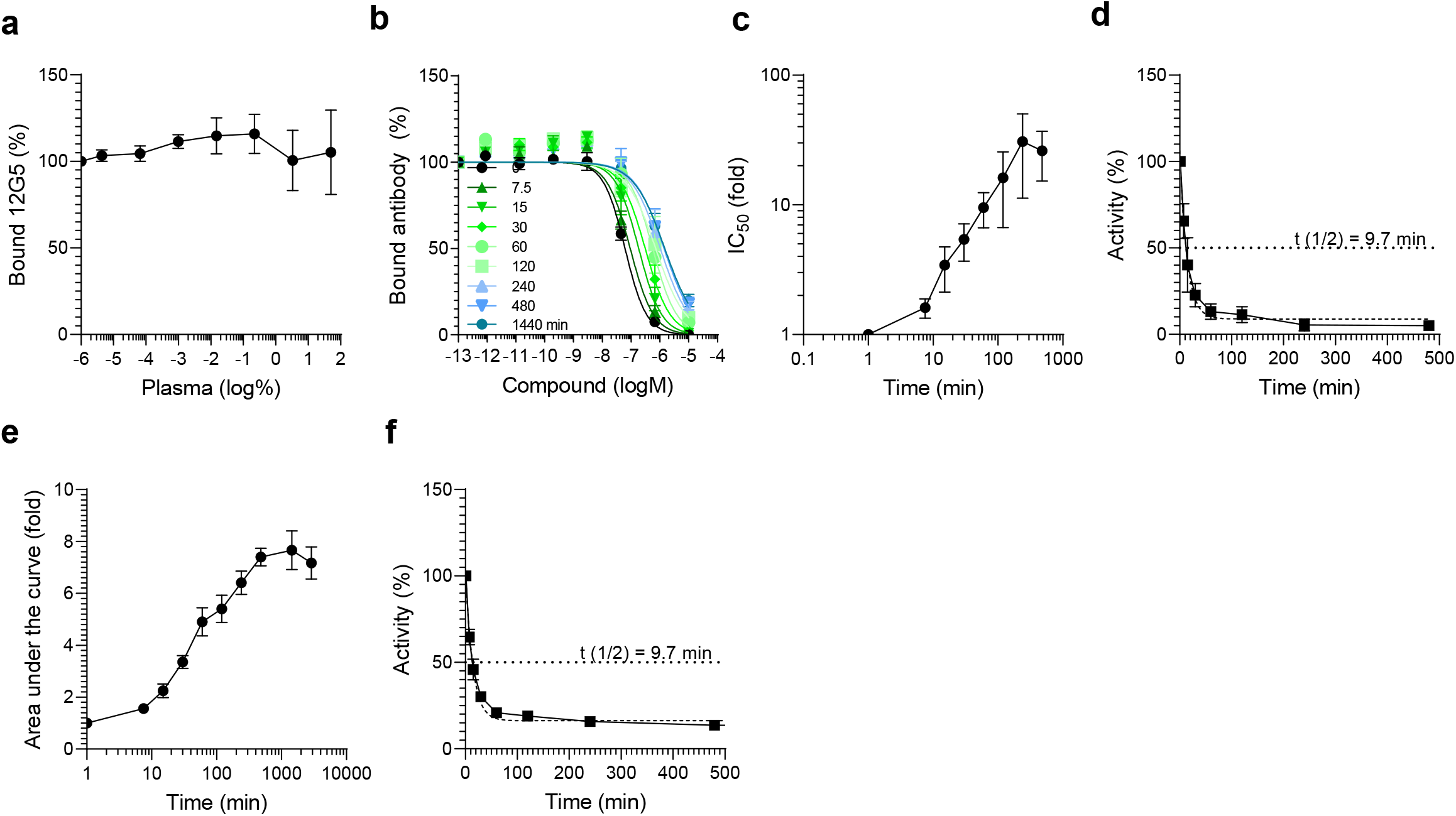
Adaption of the antibody competition assay to determine plasma stability of EPI-X4 WSC02. The peptide (or no peptide as control) was diluted in human plasma (99.3 %) at a concentration of 20 μM and incubated at 37°C for 24 h. Aliquots were taken at indicated times and stored at −80°C. Samples were defrosted simultaneously, diluted in PBS, mixed with 12G5-APC antibody and applied to the cells. Bound antibody was analyzed after 2 h of incubation at 4°C by flow cytometry. (a) Plasma alone does not affect 12G5 CXCR4 mAb binding to CXCR4. (b) Antibody inhibition curves obtained at different time points. (c) IC_50_ values obtained from curves shown in (b) were calculated in GraphPad Prism by non-linear regression and plotted over time. The IC_50_ obtained at t = 0 was set = 1. (d) Relative CXCR4 binding activity as compared to t = 0. The half-life was determined by One-phase-decay model using GraphPad Prism. (e) The area under the inhibition curves were determined from data shown in (b) using GraphPad Prism by non-linear regression. AUCs were normalized relative to t = 0. f) AUCs were transformed to obtain the activity for given time points relative to t = 0. Shown are data derived 3 individual incubation rounds using a plasma pool derived from 6 donors ± SEM.

The time point where the peptide has lost half of its antibody-blocking activity was defined as its half-life. For its determination, we calculated the fold-change of the IC_50_ values for the individual time points relative to the 0 min control (Fig. 3c). The CXCR4 binding activity before incubation was then defined as 100% and the decay in receptor-binding activity calculated and plotted over time (Fig. 3d). The half-life (t_1/2_) of WCS02 in plasma was then calculated using a one-phase decay model and revealed a value of ~ 10 min (Fig. 3d). As an alternative way to determine plasma stability, we considered the individual areas under the curve (AUCs) values rather than the IC_50_ values. Similar curve shapes and decay curves (Fig. 3e, f) and identical average half-lives were determined by both methods, validating both models. Thus, the 12G5 competition assay allows a rapid and convenient determination of peptide activity in 100% human plasma.

### Plasma stability of EPI-X4 derivatives

Next, we determined plasma stabilities of small molecules AMD3100 and IT1t or several EPI-X4 derivatives. All compounds (20 μM) were resuspended simultaneously in 100% pooled human plasma, and mixtures were incubated at 37°C. Aliquots were taken and analyzed essentially as described above. All CXCR4 antagonists obtained at t = 0 blocked 12G5 binding in a concentration-dependent manner (Fig. 4a, 4b). However, peptide-based ligands lost antibody-competing activity over time (Fig. 4b, Fig. S5) whereas AMD3100 and IT1t remained fully active, even after 24 h and, in the case of AMD3100, 48 h of incubation (Fig. 4a). The extrapolated and calculated t_1/2_ for EPI-X4 was 23 min (Fig. 4a, Fig. S5), confirming previous data obtained with an EPI-X4 specific sandwich ELISA^31,51^. The improved EPI-X4 derivative WSC02 was less stable (t_1/2_ of 10 min) (Fig. Fig. 4b, Fig. S5), perhaps because of the L1I exchange at the N-terminus^31^. However, the corresponding dimer in which two WSC02 molecules are linked via a -S-S- bridge was 19-fold more stable (t_1/2_ of 193 min) (Fig. 4b, Fig. S5), suggesting that dimerization prevents the accessibility of the N-terminal Ile residues for leucyl-aminopeptidases. We then analyzed whether storage of plasma may affect its proteolytic activity. We found that plasma that was stored at 37°C showed reduced WSC02 inactivating activity (Fig. S6). These results indicate that i) plasma which was obtained and stored under ill-defined conditions should not be used for stability assays, and ii) that meaningful data can only be produced within the first 20 h.

**Figure 4.**
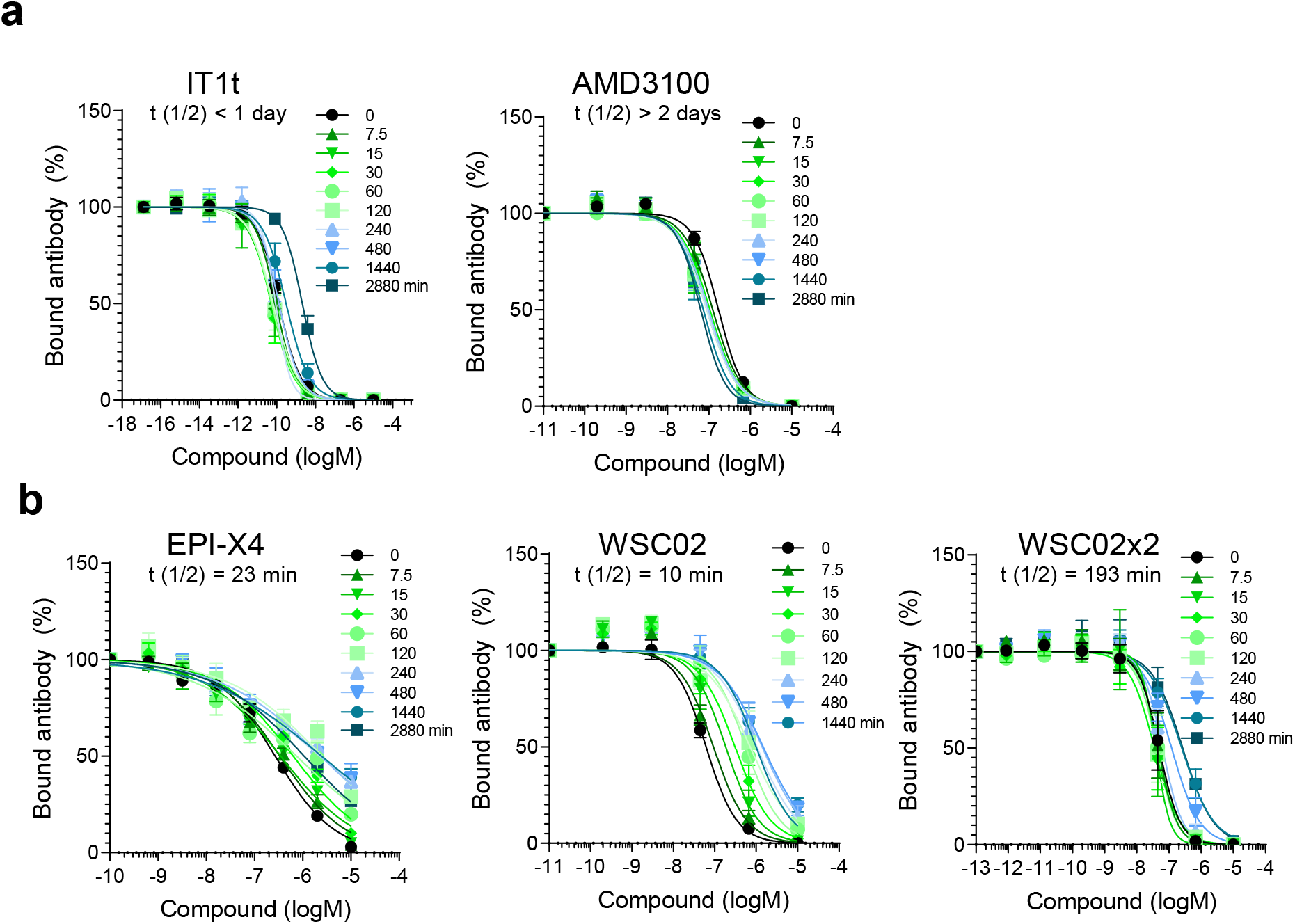
Determination of the functional half-life of CXCR4 ligands in plasma by antibody competition assay. (a) Small molecules IT1t or AMD3100 or (b) peptides EPI-X4, WSC02, and WSC02×2 were diluted in 99.3 % plasma resulting in 20 μM concentrations. Samples were incubated at 37°C, aliquots taken at indicated time points and stored at −80°C. Immediately prior to the competition assay, samples were defrosted, serially diluted in PBS, mixed with 12G5-APC antibody and applied to the cells. Bound antibody was analyzed after 2 h of incubation at 4°C by flow cytometry. The functional half-life (t_1/2_) was determined by calculating the IC_50_ values of the individual inhibition curves by non-linear regression, which are expressed relative to the t = 0 time point (100 % activity), and applying one-phase-decay model using GraphPad Prism (see Fig. S5). Shown are data derived from three independent rounds of incubation ± SEM.

Next, we evaluated whether the competition assay may be applied to measure peptide degradation in mouse plasma. EPI-X4 WSC02 was incubated in 100 % human (pooled) and mouse plasma (obtained from a single animal) for up to 60 min. Mouse plasma alone did not affect 12G5 binding at concentration of up to 50 % in cell culture, similar as human plasma (Fig. S8a). Loss of CXCR4 binding activity for EPI-X4 WSC02 was observed for the incubation in both plasma (Fig. S8b), with a more rapid inactivation of the peptide in mouse plasma (Fig. S8c), likely reflecting the increased proteolytic activity in rodents versus humans.

### Whole blood stability of CXCR4 ligands

Measuring the stability of drugs in plasma not necessarily reflects the real conditions in whole blood. To test whether the competition assay performs in blood, we used fresh EDTA blood obtained from an individual donor. To allow comparison between blood and plasma, half the of the blood sample was centrifuged to obtain plasma. The different CXCR4 ligands were diluted to 20 μM in both fluids, samples were agitated at 37°C, aliquots taken and frozen at −80°C. To check for unspecific effects, plasma and blood alone were incubated. Samples were thawed in parallel, centrifuged to get rid of cells and debris, and simultaneously analyzed for CXCR4 12G5 mAb competition. Whole blood at concentrations of up to 50 % in cell culture did not interfere with 12G5 binding (Fig. 5a), similar to plasma (Fig. 3a and 5a). As observed in the experiment before with pooled plasma, IT1t was moderately impaired in its ability to block 12G5 binding upon incubation in plasma derived from the individual donor as well as whole blood (Fig. 5b), and AMD3100 remained fully active even after 24 h (Fig. 5c). Notably, both WSC02 (Fig. 5d) and its dimer WSC02×2 (Fig. 5e) were less stable in blood compared to plasma.

**Figure 5.**
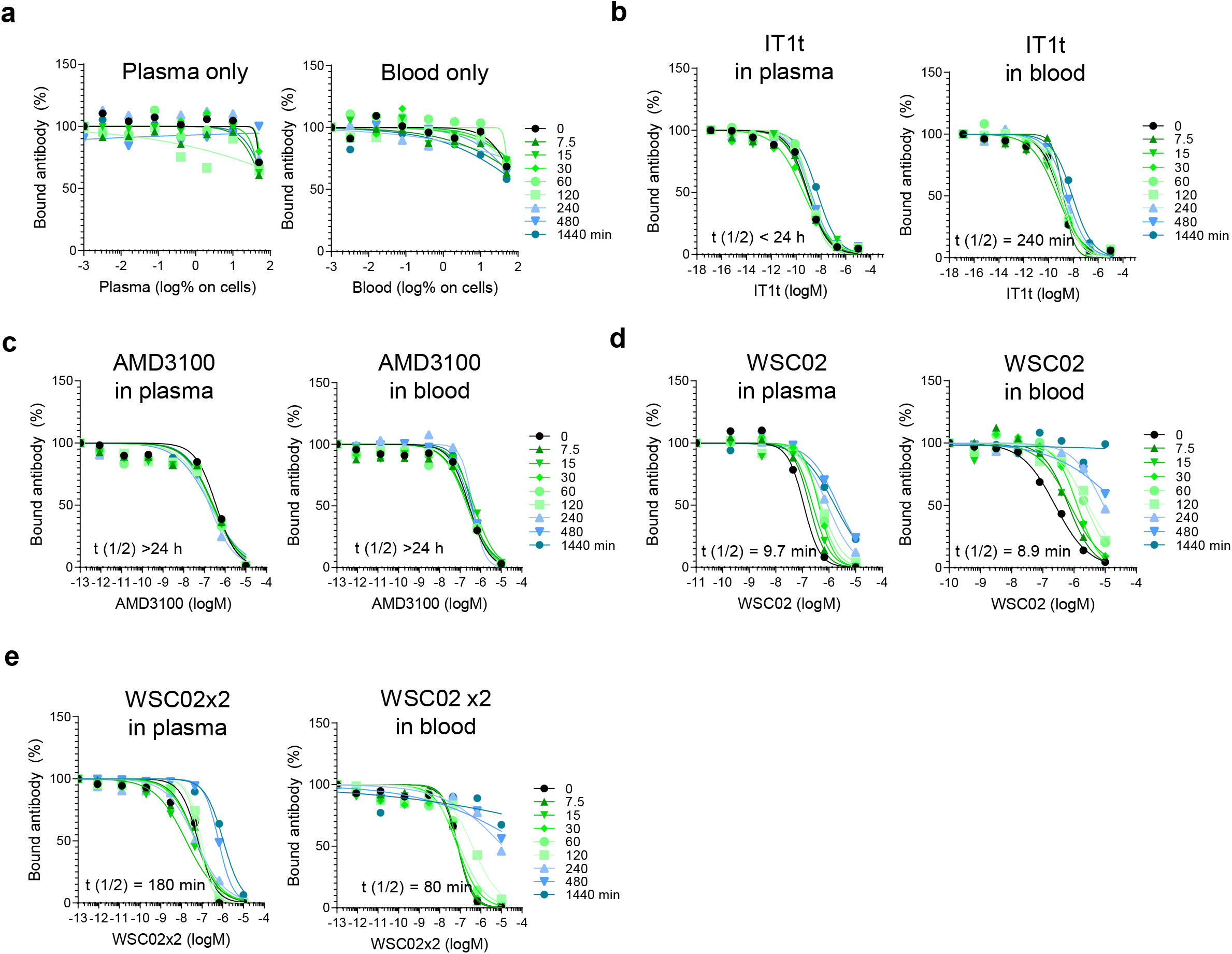
CXCR4 ligand activity and half-life in whole blood. No peptide (a), IT1t (b), AMD3100 (c), WSC02 (d) or WSC02×2 (e) were diluted in plasma or whole blood obtained from the same donor. Samples were incubated at 37°C, and aliquots taken and frozen at different time points. Samples were defrosted and centrifuged to remove cells and debris before competition with the 12G5 CXCR4 mAb of all aliquots was done simultaneously as described in Fig. 4. Percent activity was calculated by determining IC_50_ values for each time point relative to the activity at t = 0 (100%). The half-life was calculated using GraphPad Prism applying a one-phase-decay model (see Fig S. 7). Data shown are derived from an individual experiment.

## DISCUSSION

Antibody competition assays have long been used to study the interaction of ligands with receptors, e.g chemokine receptors^52,53,25,54^. There are several advantages in using antibodies instead of chemokine ligands for these kinds of assays. On the one hand, chemokines are the natural receptor ligands and therefore of high physiological relevance. On the other hand, they often have the disadvantage of being not specific for only one receptor^55^. CXCL12 for example, the natural chemokine ligand for CXCR4, also binds with high affinity to CXCR7 making it necessary to test all cells for expression of this receptor beforehand^56^. Antibodies are usually very specific for their target and can therefore be used on any desired cell type. Also, chemokines perform a two-step binding mode^57^. They first bind to the N-terminus of the GPCR and afterwards to the orthosteric binding pocket. If a small molecule blocks binding to the orthosteric binding site, it could happen that the labelled chemokine still “sticks” to the N-terminus of the receptor. This effect was, for example, demonstrated for CXCL12 that could not completely be released from CXCR4 upon binding of AMD3100^58^. This might lead to a background signal during the FACS measurement^54^. Another very important point is that chemokine binding leads to cell signaling and often to internalization. Background signal caused by internalized fluorescent ligand can be minimalized by using an antibody, that does not lead to internalization of the receptor. Also, labelled chemokines are comparably more expensive than most antibodies, what limits the throughput of such assays especially in smaller laboratories. For those reasons, we here describe a competition binding assay based on a fluorescently labelled antibody that allows the rapid determination of binding affinities of CXCR4 antagonists *in vitro* and its adaption to determine stability of ligands in human or mouse blood *ex vivo.*

A prerequisite for this assay is that ligand and antibody share orthosteric binding sites on the receptor, resulting in a competitive binding mode. Because of its importance as drug target, CXCR4 is a well-researched receptor, and a variety of antibodies targeting different epitopes on CXCR4 exist. The monoclonal antibody 12G5 binds a region in the second extracellular loop 2 (ECL2) of CXCR4, and competitively abrogates binding by the chemokine ligand CXCL12^40,41,59^. Similarly, antibody competition assays with 12G5 demonstrated that ECL2 serves as binding site for many CXCR4 antagonizing compounds that are currently in (pre)clinical development.

We here confirm these data and show that all analyzed peptides (BKT140, CTCE-9908, LY2510924, EPI-X4 and its optimized derivative) and small molecule (AMD3100, AMD070, IT1t, TG-0054) inhibitors of CXCR4 compete with 12G5 binding. The only exception was MSX-122, a biased CXCR4 ligand that intervenes in the Gα_i_-signaling pathway (cAMP modulation), but not the Gq-pathway (calcium flux)^20^. Interestingly, molecular docking experiments suggested that MSX-122 binds in close proximity to a spacious pocket in CXCR4 that is also occupied by BTK140. MSX-122 (292 Da) is the smallest of all tested compounds and binds near the bottom of the binding pocket^20^. BTK140 (2159 Da) is 7 times larger but displaced 12G5, suggesting that MSX-122 is simply too small to interfere with 12G5 binding to ECL2. In agreement with this assumption are previous studies reporting that MSX-122 is not preventing CXCL12 binding, and our data presented here that MSX-122 is not affecting CXCR4-tropic HIV-1 infection, in contrast to most other tested CXCR4 antagonists.

The fluorescence-based antibody competition assay allows for a rapid and quantitative determination of the half-maximal inhibitory concentration (IC_50_) and the inhibitory constants (K_i_) for each CXCR4 ligand analyzed, allowing a direct comparison of pharmacological parameters. The lowest K_i_, (~ 0.1 nM) was observed for Burixafor (TG-0054), an orally bioavailable stem cell mobilizing agent currently in phase I trials^23^, followed by IT1t (K_i_ ~ 0.6 nM), an allosteric inhibitor that is currently not in clinical development. AMD3100, the only marketed CXCR4 antagonist approved to mobilize hematopoietic stem cells in cancer patients^60^ and currently also in advanced clinical trials for therapy of other CXCR4-associated diseases, showed a high nanomolar K_i_ with 221 nM. For peptide-based drugs, the lowest K_i_ value was obtained for the cyclic LY251092 which is in phase I and II as part of a combination therapy against cancer. The dimeric EPI-X4 derivative WSC02×2 showed a K_i_ of 69 nM, which is ~10-fold lower than that of the endogenous peptide and reflects also its enhanced potency in inhibiting CXCR4-tropic HIV-1 infection and CXCL12-dependent cell migration. Due to the high assay performance (quick and easy) we are currently applying the 12G5 competition assay in an ongoing QSAR study with more than 200 EPI-X4 analogs in order to identify leads with reduced Ki values for further preclinical development.

The greatest advantage of the assay is to quickly and accurately determine the *functional* stability of CXCR4 ligands in plasma and whole blood by measuring the *activity* (competition with 12G5 for binding to CXCR4) rather than physical *presence* of the drugs. From the availability of the samples until the completion of the analysis by flow cytometry and evaluation of the data, it only takes 3 hours per 96 well plate. Furthermore, no sample processing is required as the peptide-containing specimen (plasma/blood) is applied directly to the cells together with the antibody. This is in sharp contrast to all other approaches, mainly mass spectrometry-based (MS) methods, that directly quantify the presence (or absence) of the analyte. These measures are time consuming and labor intensive, as the analyte containing plasma/blood sample first has to be processed to extract peptides/proteins, followed by chromatographic separation and quantitative MS, which often requires additional internal standards and expensive equipment and experienced operators. Furthermore, the antibody-based assay also allows to quantify receptor-binding of CXCR4 ligands that are bound to plasma proteins (data not shown), which may otherwise not be detected by MS. For example, 58% of administered AMD3100 directly binds to plasma proteins including albumin^61,62^ and may be missed by MS analysis.

Another advantage of the 12G5 antibody competition assay is that it enables analysis of peptide stability/activity not only in human plasma or blood but also in the respective body fluids from mice. Mouse models are normally the first choice to determine toxicity, pharmacokinetic and pharmacodynamic parameters of drugs *in vivo*, and usually involve killing the animals. Our assay precedes *in vivo* experiments and allows to identify candidates that are rapidly degraded (or even cytotoxic) prior to the animal studies. Thus, these candidate drugs will be eliminated from further analysis, which should substantially reduce animal numbers. Moreover, based on the stability profiles obtained *in vitro*, we may have first predictive values for doses to be applied later *in vivo*.

The competition assay is particularly useful when studying the stability of drug candidates that are prone to inactivation or degradation in blood, such as peptides. Our results obtained herein for the plasma half-life of the peptide inhibitor EPI-X4 (t_1/2_ 23 min) largely confirm those previously obtained using an EPI-X4 sandwich ELISA (17 min)^31^. We also corroborate that leucyl aminopeptidases are responsible for EPI-X4 degradation in plasma^31^, because addition of an inhibitor of these enzymes decreased proteolytic inactivation. The optimized derivative WSC02 was shown to have a t_1/2_ of 10 min in plasma and 9 min in blood. Interestingly, the dimeric version coupled through a disulfide bridge, WSC02×2, showed increased stability (or resistance to proteolytic inactivation) with t_1/2_ in plasma of greater than 3 h and in blood of 80 min. This finding suggests that dimerization may protect the N-terminal Isoleucine residues from efficient recognition or proteolysis by leucyl aminopeptidases. Thus, the 12G5 competition assay not only enables to determine binding affinities of EPI-X4 derivatives but also to quickly measure peptide stability in mouse and human blood, and is now routinely used as gold assay in advanced SAR studies that aim to generated highly active and stable EPI-X4 derivatives.

## METHODS

### Reagents

CTCE-9908 (#5130), and IT1t (#4596) was obtained from Tocris, Burixafor was obtained from MedKoo Biosciences (#206522), AMD070 (#HY-50101A), MSX-122 (#HY-13696), BKT-140 (#HY-P0171), and LY2510924 (#HY-12488) was obtained from from Hycultec, AMD3100 octahydrochloride hydrate was obtained from Sigma-Aldrich (#A5602). Substances were diluted at a stock concentration of 10 mM. EPI-X4 and WSC02 were synthesized automatically on a 0.10 mmol scale using standard Fmoc solid phase peptide synthesis techniques with the microwave synthesizer (Liberty blue; CEM). Amino acids were obtained from Novabiochem (Merck KGaA, Darmstadt, Germany). Peptides were purified using reverse phase preparative high-performance liquid chromatography (HPLC; Waters) in an acetonitrile/water gradient under acidic conditions on a Phenomenex C18 Luna column and afterwards lyophilized on a freeze dryer (Labconco). Prior to use peptides were diluted in PBS at a stock concentration of 3 mM. Recombinant human SDF-1α (CXCL12) was obtained from Peprotech and reconstituted at 100 μg/mL in water. WSC02-dimer (WSC02×2) was prepared by on-air-oxidation of the monomer via disulfide bonds at the cysteines in aquatic solution (pH 8.0) at 1 mg/mL peptide concentration. The dimerized peptide was then chromatographically purified and freeze dried. APC mouse anti-human CD184 (clone 12G5; #555976) and PE rat anti-human CD184 (clone 1D9; #551510) and the appropriate isotype controls were obtained from BD Pharmingen ™. L-leucinethiol (LAP) was obtained from Sigma Aldrich (#L8397).

### Blood and blood plasma

Whole blood was collected from healthy donors in EDTA tubes and directly used or subsequently centrifuged for 15 min at 2,500 x g to obtain plasma. Plasma from 6 donors was pooled and stored in aliquots at −80°C. Mouse plasma was obtained by heart punctation of BL6 mice after cervical dislocation. Blood was 19:1 diluted with 0.16 M NaEDTA and centrifuged at 2000 rcf for 20 min at 4°C to obtain plasma.

### Statement

All experiments and methods were performed in accordance with relevant guidelines and regulations. All experimental protocols were approved by a named institutional/licensing committee. Experiments involving human blood and plasma were reviewed and approved by the Institutional Review Board (i.e., the Ethics Committee of Ulm University). Informed consent was obtained from all human subjects. All human-derived samples were anonymized before use. All animal experiments were performed in accordance with the Directive 2010/63/EU of the European Parliament and approved by the regulatory authority of the state of Baden-Württemberg and in compliance with German animal protection laws.

### Cell culture

TZM-bl HIV-1 reporter cells stably expressing CD4, CXCR4 and CCR5 and harboring the lacZ reporter genes under the control of the HIV LTR promoter were obtained through the NIH AIDS Reagent Program, Division of AIDS, NIAID, NIH: TZM-bl cells (Cat#8129) from Dr. John C. Kappes, and Dr. Xiaoyun Wu. TZM-bl cells and HEK293T cells were cultured in DMEM supplemented with 10 % fetal calf serum (FCS), 100 units/mL penicillin, 100 μg/mL streptomycin, and 2 mM L-glutamine (Gibco). SupT1 cells were cultured in RPMI supplemented with 10% FCS, 100 units/mL penicillin, 100 μg/mL streptomycin, 2 mM L-glutamine and 1 mM HEPES (Gibco).

### HIV-1 inhibition assay

Viral stocks of CXCR4-tropic NL4-3 were generated by transient transfection of 293T cells with proviral DNA as described^63^. The next day the transfection mixture was removed and fresh medium containing 2.5 % FCS was added. 2 days after transfection the supernatant was harvested and cell debris were removed by centrifugation. Aliquots were stored at −80°C. For infection of TZM-bl cells in presence of inhibitors, cells were seeded at a density of 1 × 10^5^ in 70 μL DMEM containing 2.5 % FCS. Compounds were diluted in PBS and 10 μL were added. After 15 min of incubation cells were inoculated with 20 μL of diluted virus. Infection rates were determined three days after using Gal-Screen system (Applied Biosystems).

### Cell migration

Migration Assays were performed using 96-well transwell assay plates (Corning Incorporated, Kennebunk, ME, USA) with 5 μm polycarbonate filters. First, 50 μl (0.75 × 10^5^) of SupT1 cells resuspended in assay buffer (RPMI supplemented with 0.1 % BSA) were added into the upper chamber together with/without the compounds and allowed to settle down for around 15 min. In the meanwhile, 200 μl assay buffer with or without 100 ng/ml CXCL12 were filled into a 96 well-V plate. To avoid diffusion of the compounds from the upper to the lower compartment compounds were added in the same concentrations to the lower chamber. Next, the upper chamber was placed onto the 96 well-V plate and cells were allowed to migrate towards CXCL12 for 4 h at 37°C (5 % CO_2_). Afterwards, migrated cells in the lower compartment were directly analyzed by Cell-Titer-Glo® assay (Promega, Madison, WI, USA). The percentage of migrated cells was calculated as described by Balabanian et al (2005)^64^. To determine the relative migration in %, the percentage of migrated cells was normalized to the CXCL12 control without inhibitors.

### Quantitative flow cytometry

CXCR4 levels on SupT1 cells were determined using QIFIKIT® (Agilent Dako) according to the manufacturer’s protocol. Shortly, SupT1 cells were labelled with a primary CXCR4 antibody (clone 12G5) at a saturating concentration. Then the cells in parallel with provided beads coated with defined quantities of a mouse monoclonal antibody are incubated with a fluorescently labelled secondary antibody and analyzed by flow cytometry. The mean fluorescence of the cells was then interpolated with the bead’s calibration curve.

### Antibody competition assay

Competition of compounds with antibody binding was performed on SupT1 cells. For that cells were washed in PBS containing 1 % FCS by centrifugation (1.300 rpm, 4°C) and 5,000 cells (for K_i_ determinations) or 50,000 cells were then seeded per well in a 96 V-well plate. Buffer was removed by centrifugation and plates were precooled at 4°C for at least 15 min. Compounds were serially diluted in ice cold PBS and 12G5-APC antibody was diluted in cold PBS containing 1 % FCS. 15 μL of compounds was then added to the cells and 15 μL antibody immediately afterwards (0.245 nM final concentration). Plates were incubated at 4°C in the dark for 2 hours. Afterwards cells were washed twice by centrifugation with PBS containing 1 % FCS and fixed with 2 % PFA. During all pipetting steps, plates were kept cool by using a cooling pad. Antibody binding was analyzed by flow cytometry (FACS CytoFLEX; Beckman Coulter®) by determination of the mean fluorescence intensity (MFI) of at least 5.000 cells (50.000 cells per well) or 1.000 cells (5.000 cells per well).

### Formulas

*K*_*d*_ calculation for 12G5

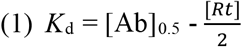

For [Ab]_0.5_ = concentration where free ligand [*L*] is *K*_*d*_

*K*_*d*_ = dissociation constant

*R*_*t*_ = total receptor concentration; so:

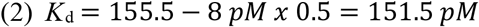

*K*_i_ calculation

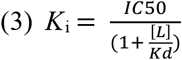

For *K*_i_ = inhibitory constant

*IC*_50_ = half maximal inhibition for 12G5 binding

[*L*] = 12G5 concentration

*K*_d_ = dissociation constant of 12G5

### Stability measurements in whole human plasma or blood

Compounds were 150-fold diluted in human plasma or whole human blood to reach final concentrations of 20 μM. The t = 0 sample was immediately taken and stored at −80°C. Plasma/compound or blood/compound mixture was then transferred to 37°C and shook at 350 rpm. At given time points samples were taken and stored at −80°C°. For measuring the functional activity of the plasma/peptide samples, the mixtures were thawed and serially diluted in ice cold PBS (starting with 100 % sample). 12G5-APC antibody competition was then performed as described before. Note, that during the competition process the samples are 1:1 diluted with the antibody and the highest plasma concentration on cells was 50 %. For blood/peptide functional stability, samples were thawed and centrifuged at 14,000 rpm for 3 min at 4°C to remove cells and debris. The supernatant was then serially diluted in PBS (starting with 100 % sample) and 12G5-antibody competition assay was performed. After the 2 h incubation, the cells were washed by centrifugation (1.300 rpm, 4°C) and 50 μL of 1-step-Fix/Lyse solution (Thermo Fisher #00-5333-54) was added for 15 min at room temperature. Afterwards cells were washed again and analyzed for bound antibody.

### Calculations and statistical analysis

Statistical analysis was performed in GraphPad Prism (version 8.3.0). IC_50_ and IC_90_ curves and the areas under the curves (AUC) were determined by nonlinear regression. For half-life calculations, the IC_50_ valuesfor each time point as well as the AUCs were analyzed and normalized relative to t = 0. Activities were then determined ((IC_50_ (t = 0) / IC_50_ (t)) × 100) for each time point and half-lives determined by a One-Phase-Decay model.

## Acknowledgements

J.M. and L.S. gratefully acknowledge the German Research Foundation (DFG) for funding the present study within the CRC1279 framework. J.M. also acknowledges funding through the Baden-Württemberg Stiftung and the European Research Council.

## Author contributions statement

M.H. conceived and conducted all experiments except for the migration assays, carried out by A.G. L.S. synthesized and analyzed peptides. A.B. and V.R. helped with all mice experiments. C.G. and B.M. interpreted data and assisted in writing the manuscript. J.M. supervised the study and wrote the paper together with M.H.

## Additional Information

Supplementary information accompanies this paper at …,.

