## Supplement for "Microtiter plate-based antibody-competition assay to determine binding affinities and plasma/blood stability of CXCR4 ligands"

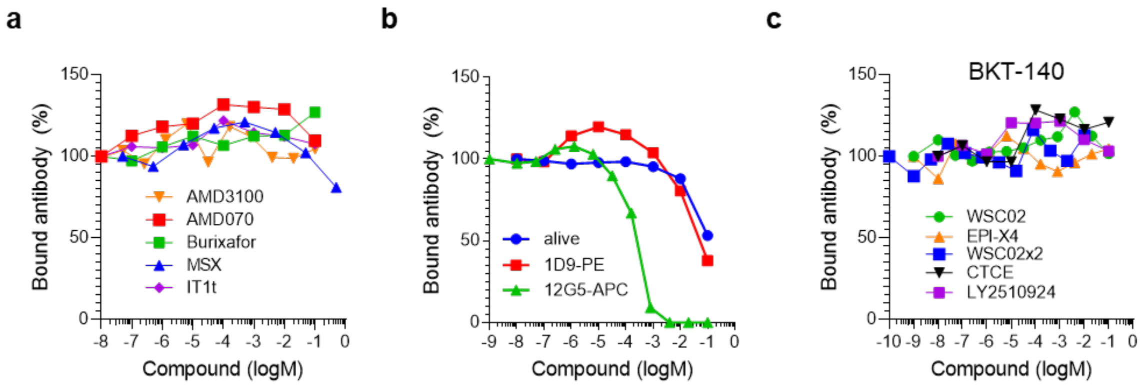


**Figure S1. CXCR4 ligands do not compete with control-antibody binding (1D9)**. Small molecule (a) or peptide (b, c) ligands were diluted in PBS and added to precooled SupT1 cells. A constant concentration of PE-labelled 1D9-antibody (or 12G5-APC antibody in (c)) was added immediately afterwards. After 2 h incubation in the dark, cells were washed and analyzed by flow cytometry. (c) For BKT-140, percentage of viable cells was determined by measuring the percentage of gated living cells in flow cytometry upon treatment with the inhibitor. Shown are data derived from a single experiment.

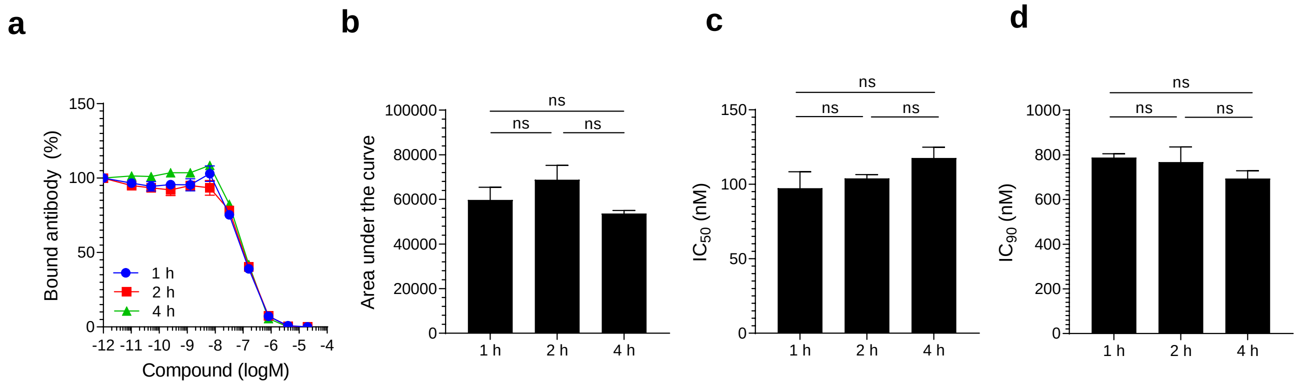


**Figure S2. 12G5 antibody binding and competition at various timepoints.** EPI-X4 WSC02 was titrated on SupT1 cells in presence of a constant concentration of 12G5-APC CXCR4 antibody and incubated at 4°C for 1, 2 and 4 h. Afterwards unbound antibody was removed and cells were analyzed by flow cytometry. (a) Antibody-inhibition curves for all time points. (b) Area under the curves (AUC) obtained from a; (c) $IC_{50}$ values and (d) $IC_{90}$ values were determined from data shown in (a) using GraphPad Prism by non-linear regression. Shown are average values derived from one experiment performed in triplicates ± SEM. ns = non-significant (one way-ANOVA).

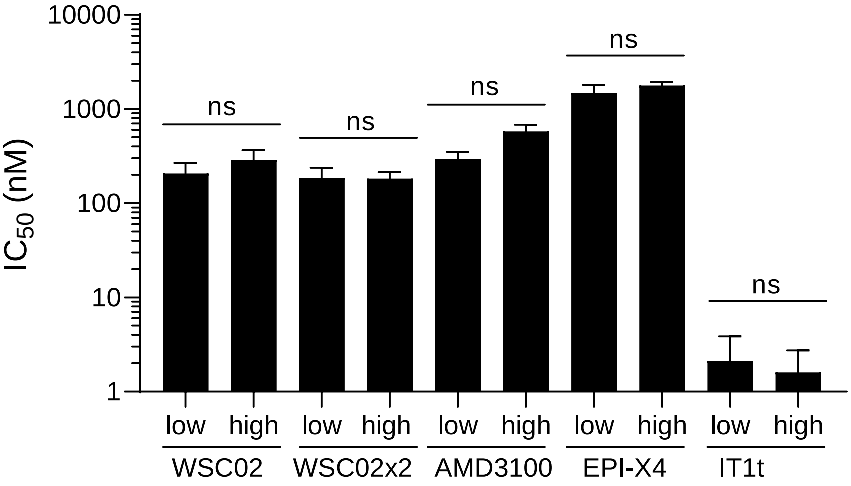


**Figure S3. A high or low cell number does not affect $IC_{50}$ values obtained in the competition assay.** The antibody competition assay was performed in the presence of 5,000 (low) or 50,000 cells (high). $IC_{50}$ values were determined using GraphPad Prism and compared by one-way ANOVA. Shown are data derived from 3 (high) or 5 (low) independent biological replicates ± SEM. ns = non-significant.

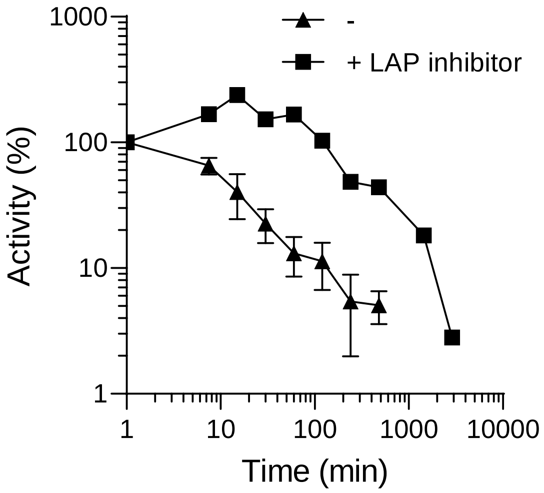


**Figure S4. Inhibition of EPI-X4 WSC02 degradation in plasma by a leucyl-aminopeptidase-inhibitor.** 20 µM of WSC02 was incubated in human plasma alone or plasma pretreated with 300 µM of L-leucinethiol (an inhibitor of leucyl-aminopeptidases (LAP)) for 10 min. At indicated time points aliquots were taken and stored at -80°C. Activity of plasma/peptide was determined using 12G5-antibody competition assay. Shown are data derived from 1 (inhibited) or 3 (no inhibitor) independent rounds of incubation ± SEM.

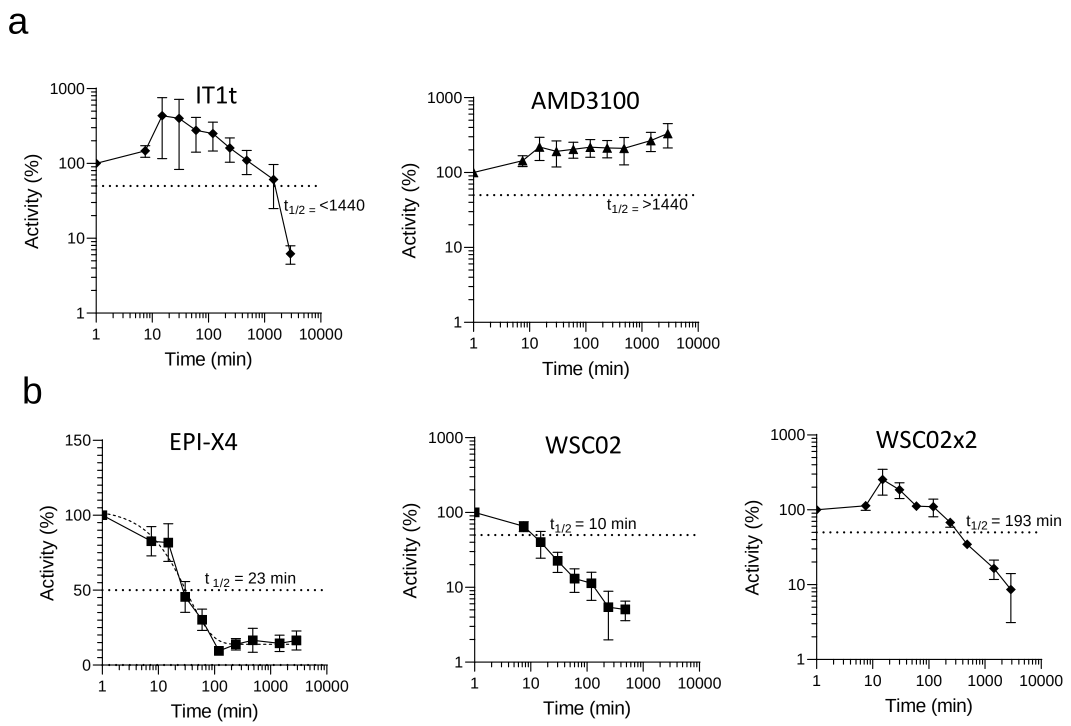


**Figure S5. Loss of CXCR4 antibody competition over time and calculation of functional half-lives.** The functional half-life ($t_{1/2}$) of small molecules (a) and peptide agents (b) was determined by calculating the $IC_{50}$ values of the individual inhibition curves shown in main Figure 4. The $IC_{50}$ values obtained at t = 0 were then defined as fully active (100 %), and the increase in the $IC_{50}$ (decrease in activity) over time was determined by calculating ($IC_{50}$ (t = 0)/$IC_{50}$ (t = x)) x 100 . The half-lives ($t_{1/2}$) were then calculated applying one-phase-decay model using GraphPad Prism (if possible) or estimated by eye. Shown are data derived from 3 - 4 individual rounds of incubation in plasma ± SEM. Also see Figure 4.

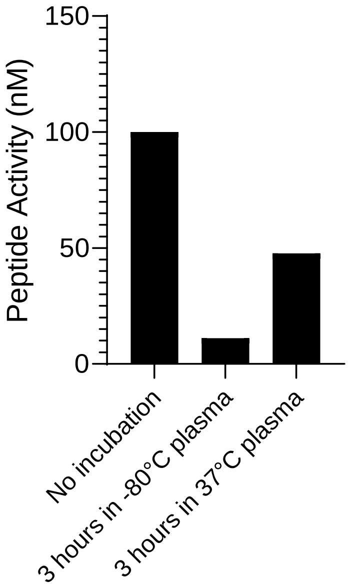


**Figure S6. Incubation of plasma at 37°C decreases its degradation-activity.** 20 µM of WSC02 was incubated in human plasma that was incubated at 37°C for 24 h. At given t = 0 and t = 3 h an aliquot was taken and stored at -80°C. Activity of plasma/peptide was determined using 12G5-antibody competition assay as described before. Data are derived from a single round of plasma incubation.

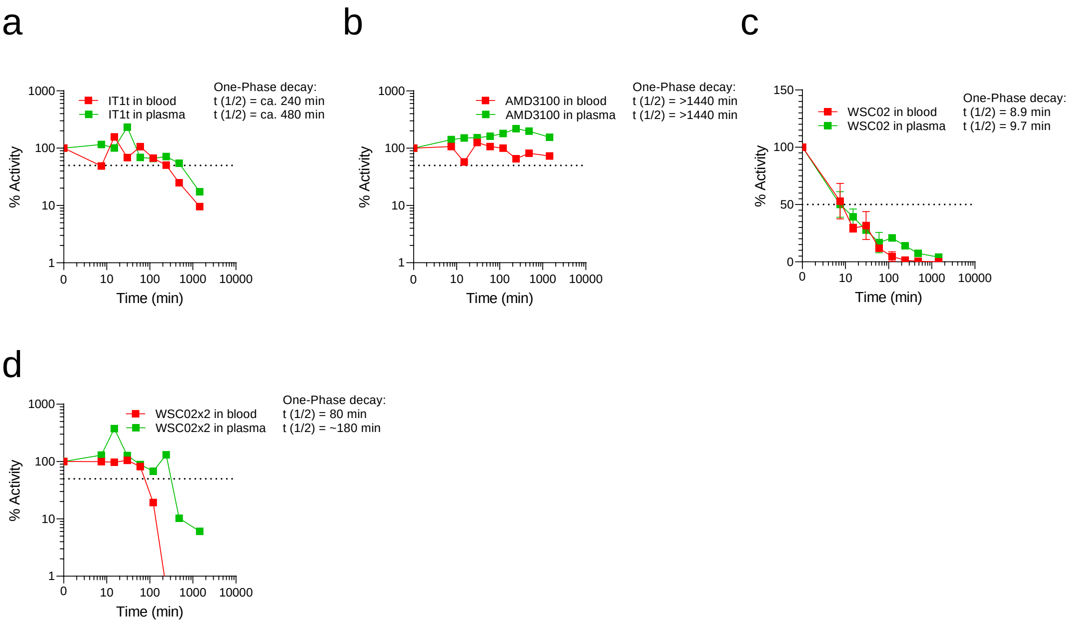


**Figure S7: CXCR4 ligand activity and half-life determination in whole blood.** IT1T (a), AMD3100 (b), WSC02 (c) or WSC02x2 (d) were diluted in plasma or whole blood obtained from the same donor. Samples were incubated at 37°C, and aliquots taken and frozen at different time points. Samples were defrosted and centrifuged to remove cells and debris before competition with the 12G5 CXCR4 mAb of all aliquots was done simultaneously as described in Fig. 4. Percent activity was calculated by determining $IC_{50}$ values for each time point by non-linear regression (see also Fig. 5) relative to the activity at t = 0 (100%). The half-life was calculated using GraphPad Prism applying a one-phase-decay model (see Fig S7). Data shown are derived from an individual experiment, or 2 individual experiments ± SEM for WSC02.

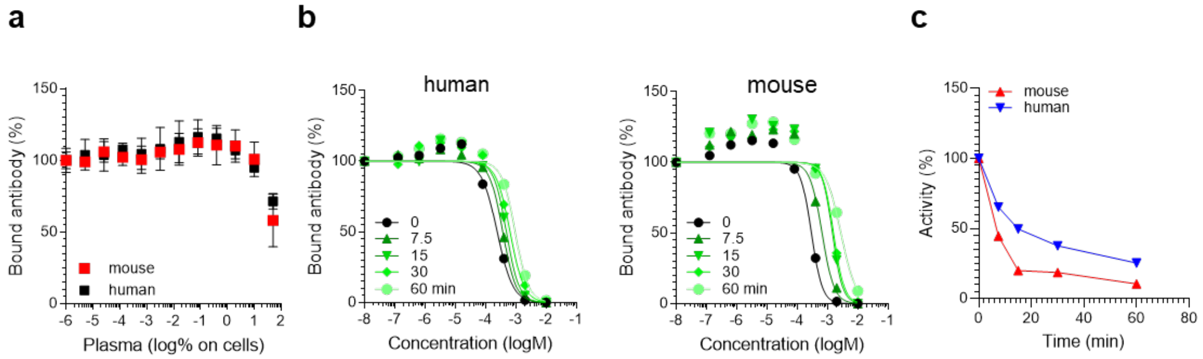


**Figure S8: Determining CXCR4 binding activity and stability of CXCR4 ligands in mouse plasma.** 20 µM of WSC02 (or mock as control) was incubated in human plasma (pool of 6 donors) or mouse plasma (single donor) for 60 min at 37°C. At given time points aliquots were taken and stored at -80°C. Activity of plasma/peptide was determined using 12G5-antibody competition assay. (a) Mouse and human plasma do not interfere with 12G5 binding at cell culture concentrations of up 50 %. (b, c) Antibody competition curves of peptides in human (left) and mouse (right) plasma. (c) From curves in (b) $IC_{50}$ values were calculated by non-linear regression and normalized relative to t = 0 and subsequently plasma half-life was calculated. Data are derived from one experiment.

**Table S1**. Parameters determined to calculate the $K_i$

| Parameter | Result | Standard deviation | n |
|---|---|---|---|
| $[Ab]_{0.5}$ of 12G5-APC | 155.5 pM | 68.2 pM | 4 |
| CXCR4 (n) on a SupT1 cell | 29,022 | 5,245 | 4 |
| Volume per well | 30 µl | - | - |
| Cells per well | 5,000 | - | - |
| CXCR4 concentration per well (Rt) | 8 pM | 1.45 pM | - |
| $K_d$ of 12G5-APC | 151.5 pM | - | - |
